# Convergently-evolved honeypot ants show mixed signs of niche divergence

**DOI:** 10.64898/2026.04.07.717096

**Authors:** Bianca Raissa Nogueira, Omar Daniel Leon-Alvarado, Lily Khadempour

## Abstract

1. Honeypot ants are a compelling system for studying convergent evolution. In these ants, specialized workers store liquid food in distended crops, a trait known as repletism, which has evolved independently across multiple genera worldwide. Seasonal resource scarcity and aridity are hypothesized drivers of this trait, yet the role of environmental factors in shaping the evolution of repletism has never been empirically tested.
2. We modeled suitable habitats for 49 honeypot ant species using ensemble distribution models, then used the most important environmental predictors to compare niche structure across and within biogeographic regions (Nearctic, Australasia, Afrotropic, Neotropic, Palearctic). To address uneven sampling density among species, we independently applied occurrence-based and Boolean (suitable-pixel) approaches, comparing each against environmental background using PERMANOVA and PERMADISP. Niche overlap was also assessed using Schoener’s D.
3. Variable importance profiles showed no consistent patterns across genera or regions, but we observed clear distinction on species occupied environmental space both within and across biogeographic regions. The relatively few cases of niche similarity and overlap occurred mainly among congeneric species and between Nearctic and Afrotropical taxa. Globally, species distributions were structured along a gradient opposing dry, thermally extreme environments and wetter, more vegetated conditions. While accumulated heat, climatic seasonality, and water-energy balance also contributed to niche differentiation.
4. Despite honeypot ants being considered desert specialists, our results show considerable environmental heterogeneity among species. As a group, honeypot ants neither occupy the same environmental space nor experience the same contemporary climatic pressures. This suggests that repletism is not a response to a single, conserved environmental condition, but to an interaction of physiological, morphological and behavioral constraints under different climatic contexts.
5. By characterizing the contemporary environmental niches of honeypot ants, this study establishes a baseline for further research on the evolution of repletism. While our results suggest that contemporary environmental factors cannot explain the convergence of honeypot ants, future work should examine paleoclimatic conditions along with species-level ecological traits.

## Abstract (in portuguese)

1. As formigas-pote-de-mel representam um sistema modelo para o estudo da evolução convergente. Nessas formigas, operárias especializadas armazenam alimento líquido em seus gásteres distendidos, uma característica conhecida como repletismo, que evoluiu independentemente em diversos gêneros ao redor do mundo. A escassez sazonal de recursos e a aridez são frequentemente apontadas como possíveis forças seletivas associadas a essa característica, mas pouco se sabe sobre o papel dos fatores ambientais na evolução do repletismo.
2. Com isso em mente, modelamos os habitats favoráveis para 49 espécies de potes-de-mel utilizando ensemble models. A partir dos modelos, os preditores ambientais mais importantes foram usados para comparações de estrutura de nicho inter e intra-regiões biogeográficas (Neártico, Australásia, Afrotropico, Neotropico e Paleártico). Para reduzir os efeitos da desigualdade na densidade amostral entre as espécies, aplicamos de forma independente abordagens baseadas em ocorrências e em mapas booleanos (pixels adequados), comparando cada uma ao background ambiental por meio de PERMANOVA e PERMADISP. A sobreposição de nicho também foi avaliada utilizando os índices D de Schoener.
3. observadas diferenças significativas no espaço ambiental ocupado pelas espécies, tanto dentro quanto entre as regiões biogeográficas. Os poucos casos de similaridade e sobreposição de nicho ocorreram principalmente entre espécies do mesmo gênero e entre táxons neárticos e afrotropicais. Em escala global, a distribuição das espécies foi estruturada ao longo de um gradiente que contrapõe ambientes secos e termicamente extremos a condições mais úmidas e vegetadas.
4. Embora as postes-de-mel sejam frequentemente consideradas especialistas de desertos, nossos resultados revelam uma grande heterogeneidade ambiental entre as espécies. Como grupo, essas formigas não ocupam o mesmo espaço ambiental e tampouco as mesmas pressões climáticas contemporâneas. Isso sugere que o repletismo não é uma resposta a uma única condição ambiental conservada, mas provavelmente é resultado da interação entre características fisiológicas, morfológicas e comportamentais sob diferentes contextos climáticos.
5. Ao caracterizar os nichos ambientais das formigas-pote, este estudo estabelece uma base importante para futuras pesquisas sobre a evolução do repletismo. Estudos futuros devem investigar condições paleoclimáticas em conjunto com características ecológicas específicas de cada espécie.

## 1 INTRODUCTION

Honeypot ants (Formicidae) display an extreme nutrient storage strategy critical for colony survival (Conway, 1986; Nogueira et al., 2026). These ants possess specialized workers called repletes that store large amounts of sugar- or protein-rich fluids in their crop for periods that can exceed one year (Hasegawa, 1993; Khalife & Peeters, 2020; Wheeler, 1915). Their gaster becomes so enlarged that gut contents are visible through their exoskeleton (Froggatt, 1896; Sawh et al., 2022; Wheeler, 1915; Figure 1), and mobility is highly compromised in most species (Snelling, 1976). Non-specialized workers bring food to repletes, which they then store. Usually, repletes are fed with sugary liquids such as nectar and honeydew (Conway, 1986; Meurville et al., 2025; Snelling, 1976). When other colony members require nutrients, repletes regurgitate and share their stored fluids through a process called trophallaxis (LeBoeuf, 2017).

**Figure 1.**
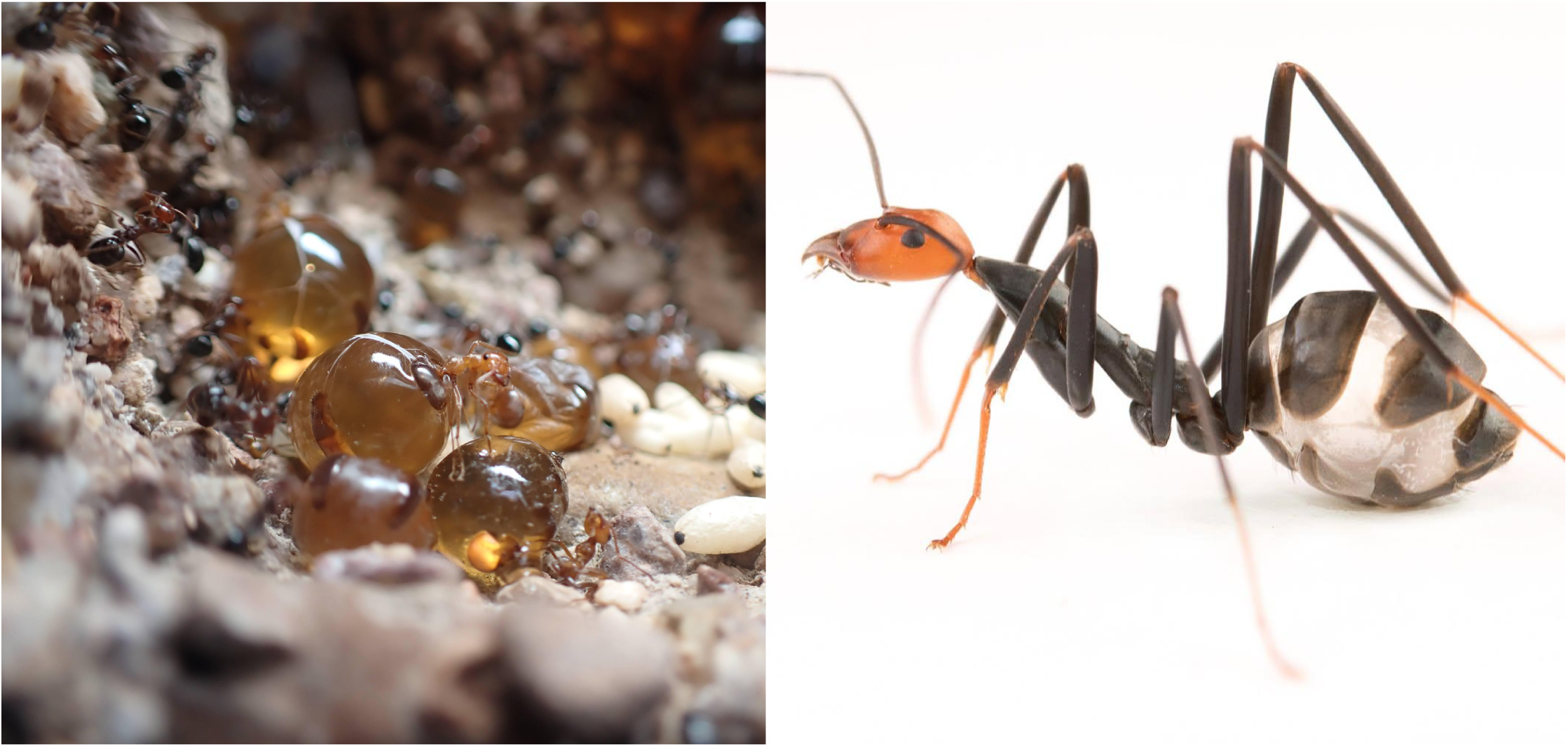
At opposite ends of the repletism spectrum, *Myrmecocystus* (left) repletes are highly distended and largely immobile, serving exclusively as living food storage, while *Leptomyrmex* (right) repletes remain mobile and continue to participate in foraging and other colony activities. Photos: *Myrmecocystus* by MP Meurville and *Leptomyrmex* by Jordan Dean.

Repletism has evolved independently in at least eight genera, primarily within Formicinae subfamily, making it a compelling example of convergent evolution (Nogueira et al., 2026; Sawh et al., 2023). Honeypot ants are found across the Neotropic, Australasia, Afrotropic, Nearctic, and Palearctic regions. *Myrmecocystus* Wesmael, 1838 is the only genus in North America and *Proformica* Ruzsky, 1902 the only one in Europe. *Leptomyrmex* Mayr, 1862, the only Dolichoderinae honeypot and the most geographically widespread genus, occurs in South America, Australia, and Melanesia. In Australia, *Melophorus bagoti* Lubbock, 1883 and *Camponotus inflatus* Lubbock, 1880 have overlapping ranges in the north and south, respectively. *Carebara perpusilla* (Emery, 1895), the only Myrmicinae known genus, and *Tapinolepis trimenii* (Forel, 1895) are distributed across southern and central Africa. In South America, *Brachymyrmex giardi* Emery, 1895 occurs in coastal Chile and *L. relictus* Boudinot & al., 2016 in central Brazil. While repletism characterizes all species in some genera (e.g., *Leptomyrmex*, *Myrmecocystus*), in others it is expressed by only a subset. *Myrmecocystus*, *Melophorus*, and *Camponotus* represent the traditional form of repletism, while remaining genera show more variable physiology, morphology, and behavior (Conway, 1986; Plowman, 1981; Wheeler, 1908, 1915).

Similar environmental pressures are hypothesized to have driven the convergent evolution of repletism (Conway, 1986). Specifically, it is thought that this trait evolved as a response to a hot, dry climate, marked by periods of resource abundance followed by prolonged scarcity. Conditions under which the ability to store food and water during periods of plenty would confer an adaptive advantage. Environmental factors that could potentially reflect these conditions are wide temperature amplitude, high seasonality, or in other words, diurnal and seasonal temperature fluctuations, and low environmental humidity (Fischer et al., 2022; Liu et al., 2026; Uhl et al., 2022).

Ecological and climatic variation, at both large and small geographical scales, play a significant role in shaping regionally-adapted phenotypes (Huey et al., 2000). Populations in harsh, climate-stressed environments can converge on functional traits essential for survival in such environments (Love et al., 2023). However, climate is not always the sole driver of convergence. Organisms in markedly different environments can show striking similarities (Alvarado-Cárdenas et al., 2013), and similar traits can arise coincidentally or as responses to past rather than current conditions (Losos, 2011; Revell et al., 2007). With this in mind, the environmental niche of honeypot ants has never been formally characterized, and the assumption that aridity drives repletism has never been tested. It remains unknown whether honeypot ants across different regions actually share environmental conditions, or whether convergence reflects a shared climatic pressure at all.

Ecological Niche Models (ENMs) represent a robust methodology for identifying suitable areas for species within both geographic and environmental spaces (Peterson et al., 2011). A defining characteristic of ENMs is their reliance on abiotic variables and the accessible area (M) of a species (Soberón & Peterson, 2005). This allows for the identification of environmental drivers that most significantly influence species contemporary distributions (Guisan & Thuiller, 2005). These techniques have been widely applied across biological disciplines, including conservation biology (Guisan et al., 2013), species delimitation (Raxworthy et al., 2007), paleo-ecology (Nogués-Bravo, 2009), and the forecasting of future distributions (Thuiller et al., 2005). In the context of honeypot ants, ENMs facilitate a comparative analysis of the variables shaping their global distribution and enable the quantification of niche overlap and differentiation within a standardized environmental space (Broennimann et al., 2012)

In this study, we aimed to determine whether honeypot ants share environmental similarities based on their current distributions. We hypothesize that honeypot ant species occupy environmental space similarly and that a similar set of environmental variables contribute to their current distribution. This hypothesis builds on the assumption that abiotic factors regulating dry habitats, where resource availability fluctuates, would make the ability to store and regulate nutrients an adaptive advantage. This study provides a comprehensive framework for understanding how environmental drivers shape contemporary species distributions. Furthermore, these findings establish a foundation for future research into how such factors influence the phenotypic evolution of honey pot ants on a global scale.

## 2 MATERIALS AND METHODS

### 2.1 Honeypot ant occurrence data

We retrieved occurrence data for 55 known honeypot ant extant species (Nogueira et al., 2026). We built a global dataset by combining records from the Global Biodiversity Information Facility (GBIF.org, 2026), published studies, and our own field collections (Derived dataset, 2026). We validated species distributions against AntMaps, which compiles published literature to map ant records globally (Guénard et al., 2017; Janicki et al., 2016). Coordinates were cleaned using the CoordinateCleaner R package (Zizka et al., 2019), and records falling at sea were reassigned to the nearest coastline within a 3 km buffer to account for projection and GPS error bias (Baselga & Olsen, 2021; Smith et al., 2023). To reduce sampling bias, we limited occurrences to one point per 1 km raster pixel (Boria et al., 2014; Hijmans, 2012; Ten Caten & Dallas, 2023). Species with fewer than five valid records after cleaning were excluded, yielding a final dataset of 4786 occurrence records across 49 species (Table S1). Although larger occurrence datasets generally improve Species Distribution Model (SDM) performance (Wisz et al., 2008), we prioritized broad species representation given the comparative nature of our study.

### 2.2 Environmental data

We assembled environmental predictors from three sources: 37 atmospheric variables from CHELSA v2.1 (Brun et al., 2022; Karger et al., 2017), including 19 bioclimatic variables and additional variables capturing temperature accumulation, growing season dynamics, atmospheric moisture, and water-energy balance; 10 soil bioclimatic variables (0-5 cm) from SoilTemp (Lembrechts et al., 2020); and percent tree and non-tree cover from MODIS Vegetation Continuous Fields (DiMiceli, et al., 2025; 2000-2024). All data were at 1 km resolution. This variable selection reflects the hypothesis that repletism evolved under seasonal resource scarcity, with stored fluids largely derived from plant nectar.

We assessed multicollinearity using VIF analysis (threshold = 8; vifstep, usdm package; Naimi et al., 2014; Zuur et al., 2010), applied sequentially within variable groups (CHELSA bioclimatic, CHELSA climate, SoilTemp) before a final combined analysis including vegetation cover. Three additional redundant variables were manually removed (ngd5, ngd10, pet_penman_range), yielding a final set of 15 predictors (Figure 2; Table S2).

**Figure 2.**
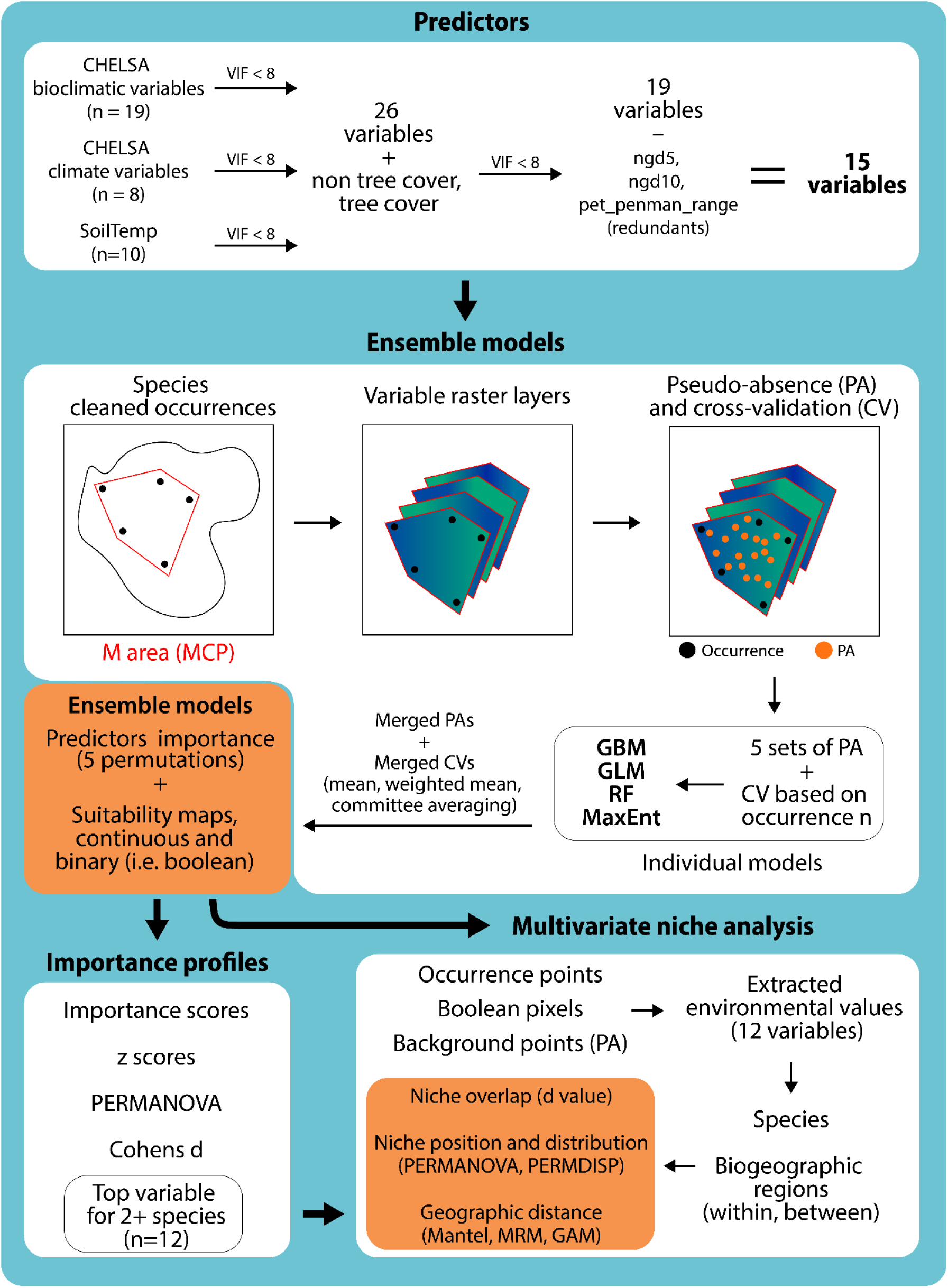
Workflow for species distribution modeling and niche analysis of honeypot ants. After VIF-based predictor filtering (VIF < 8), ensemble models were fitted per species using occurrence records, pseudo-absence sets, and cross-validation within species-specific accessible areas. Models produced suitability maps, richness maps, and variable importance scores. Environmental values extracted from both occurrence points and Boolean (suitable-pixel) data were then used in multivariate niche analyses, including PERMANOVA, PERMDISP, and pairwise niche overlap, within and across biogeographic regions.

### 2.3 Species ensemble models

We defined each species’ calibration area (M) using a Minimum Convex Polygon (MCP) with a 2.5° buffer, generated with the adehabitatHR package (v. 0.4.22; Calenge, 2006). Ensemble models (EMs) were built using biomod2 (v. 4.3-4; Thuiller et al., 2009) with four algorithms: Generalized Linear Models (GLM; Nelder & Wedderburn, 1972), Generalized Boosted Models (GBM; Elith et al., 2008), Random Forest (Prasad et al., 2006), and MaxEnt (Phillips & Dudík, 2008; Phillips et al., 2006).

We generated five pseudo-absence sets using random sampling, with the number of pseudo-absences scaled to occurrence sample size (10× occurrences; min: 1000, max: 15000) to maintain consistent presence-to-background ratios across species with highly variable occurrence densities (5-950 records; Barbet-Massin et al., 2012). Cross-validation strategy was likewise sample-size dependent: jackknife for fewer than 10 occurrences, 5-fold for 11-60, and 10-fold for more than 60, applied across all pseudo-absence sets and algorithms with a 70/30 calibration/testing split (Araújo et al., 2005). Model parameters followed the biomod2 “bigboss” strategy.

Models were evaluated using the Area Under the Precision-Recall Gain curve (AUCprg; Sofaer et al., 2019) and the True Skill Statistic (TSS; Allouche et al., 2006). We computed ensemble mean (EMmean) and weighted mean (EMwmean) across all models, with EMwmean weights proportional to individual model evaluation scores. Variable importance was assessed with five permutations per variable. Continuous suitability maps were projected onto each species’ M area and converted to binary presence/absence maps (Boolean maps) using a 10th percentile training presence threshold (P10; Pearson et al., 2007). Richness maps were generated for the most speciose regions, Australasia and the Neotropic, by projecting EMs onto occurring and adjacent ecoregions (Dinerstein et al., 2017).

#### 2.3.1 Predictor’s importance

Variable importance values were z-score standardized, and species and variables were clustered using Pearson correlation distances. We used PERMANOVA (adonis2; Anderson, 2001) with a Pearson distance matrix and 999 permutations to evaluate importance profiles across genera and biogeographic region (i.e., Neotropic, Australasia, Afrotropic, Nearctic, and Palearctic), and PERMDISP (betadisper + permutest; Anderson, 2006) to assess within-group dispersion. For significant pairwise differences, we quantified effect sizes using Cohen’s d (Cohen, 1988; Nakagawa & Cuthill, 2007) coupled with Wilcoxon tests, interpreted following established thresholds: very small (0.01), small (0.2), medium (0.5), large (0.8), very large (1.2), and huge (2.0; Sawilowsky, 2009). Mantel and partial Mantel tests (Mantel, 1967; Mantel & Valand, 1970) were used to assess whether importance patterns were associated with geographic distance.

### 2.4 Multivariate niche analysis

#### 2.4.1. Species environmental space

To evaluate whether honeypot ant species occupy a non-random subset of their available environmental space, we conducted species-level multivariate analyses using 11 environmental variables selected as important predictors for at least two species. For each species, environmental values were extracted from three sources: occurrence points, background points sampled within the accessible area (M area), and Boolean pixels from EMs predictions. A Principal Component Analysis (PCA) calibrated on background values represented the available environmental conditions, into which occurrence points and Boolean pixels were independently projected. Principal components explaining at least 80% of variance were retained for multivariate analyses of distributional differences (PERMANOVA) and dispersion (PERMDISP).

We performed three pairwise comparisons per species: occurrences vs. background, Boolean pixels vs. background, and occurrences vs. Boolean pixels. The first two tested whether occupied and predicted environmental spaces were restricted relative to what was available; the third assessed consistency between predicted and observed environmental conditions. To account for unequal sample sizes, all comparisons were repeated across 999 iterations with random subsampling to match dataset sizes (minimum 1000 points for Boolean and background comparisons). Mean and median statistics were calculated across iterations to ensure robust results.

#### 2.4.2 Niche comparisons within and between biogeographic regions

To assess whether honeypot ant species partition or share environmental space, we performed niche comparisons both within and across biogeographic regions. For within-region analyses, occurrence points and Boolean pixels were projected into regional PCA spaces calibrated with local background values, so each PCA reflects environmental variation available within that region. For between-region analyses, all 49 species were projected into a single global PCA calibrated with background values from all regions combined, placing species into a common environmental reference. In both cases, we retained the minimum number of PCs explaining 80-90% of variance.

We assessed pairwise niche differences using PERMANOVA with Benjamini-Hochberg (BH) adjusted p-values, and niche overlap using Schoener’s D calculated from two-dimensional kernel density estimates (MASS::kde2d, 100×100 grid; Venables & Ripley, 2002), with density grids normalized before computing overlap (Broennimann et al., 2012; Warren et al., 2008). Schoener’s D ranges from 0 (no overlap) to 1 (complete overlap; Schoener 1968), and values were classified as negligible (0-0.2), low (0.2-0.4), moderate (0.4-0.6), high (0.6-0.8), or near-complete (0.8-1.0).

For the between-region analyses, we calculated environmental and geographic centroids per species, and pairwise environmental distances were computed as Euclidean distances between centroids. We computed geographic distances as Haversine distances in kilometers. We tested whether geographic distance predicted environmental differentiation using a Mantel test (Spearman correlation, 999 permutations), multiple regression on distance matrices (MRM) with and without region membership, and a generalized additive model (GAM; Hastie & Tibshirani, 1986; Wood, 2017) to assess non-linearity. Overall differences among regions were tested with PERMANOVA and PERMDISP. Because only Australasia and the Nearctic had more than two co-occurring species, Mantel and GAM analyses were restricted to these regions; remaining regions were compared descriptively.

All analyses described in this section were conducted in R 4.4.1 and R 4.4.6 (R Core Team 2024) using RStudio 2024.09.0, with spatial data processing performed in QGIS 3.36.0 (Graser et al., 2025).

## 3 RESULTS

### 3.1 Honeypot ant occurrence data

After coordinate cleaning, occurrence records varied substantially among genera (Figure 3). *Leptomyrmex* was the best-represented group, with over 2700 records across 18 species, primarily in tropical and subtropical biomes. *Myrmecocystus* comprised 27 species and 1783 occurrences, mostly associated with desert environments but extending into mediterranean, temperate, and subtropical regions. Remaining genera had fewer records: *Proformica nasuta* (Nylander, 1856) (n = 70) was restricted to mediterranean environments, while *Proformica epinotalis* Kuznetsov-Ugamsky, 1927 (n = 25) inhabited temperate and marginal desert areas; *Carebara perpusilla* (n = 29), *Melophorus bagoti* (n = 62), and *Camponotus inflatus* (n = 21) were primarily associated with desert biomes; and *Tapinolepis trimenii* (n = 7) and *Brachymyrmex giardi* (n = 6) showed the most limited distributions, in desert and mediterranean biomes respectively (Figure S1).

**Figure 3.**
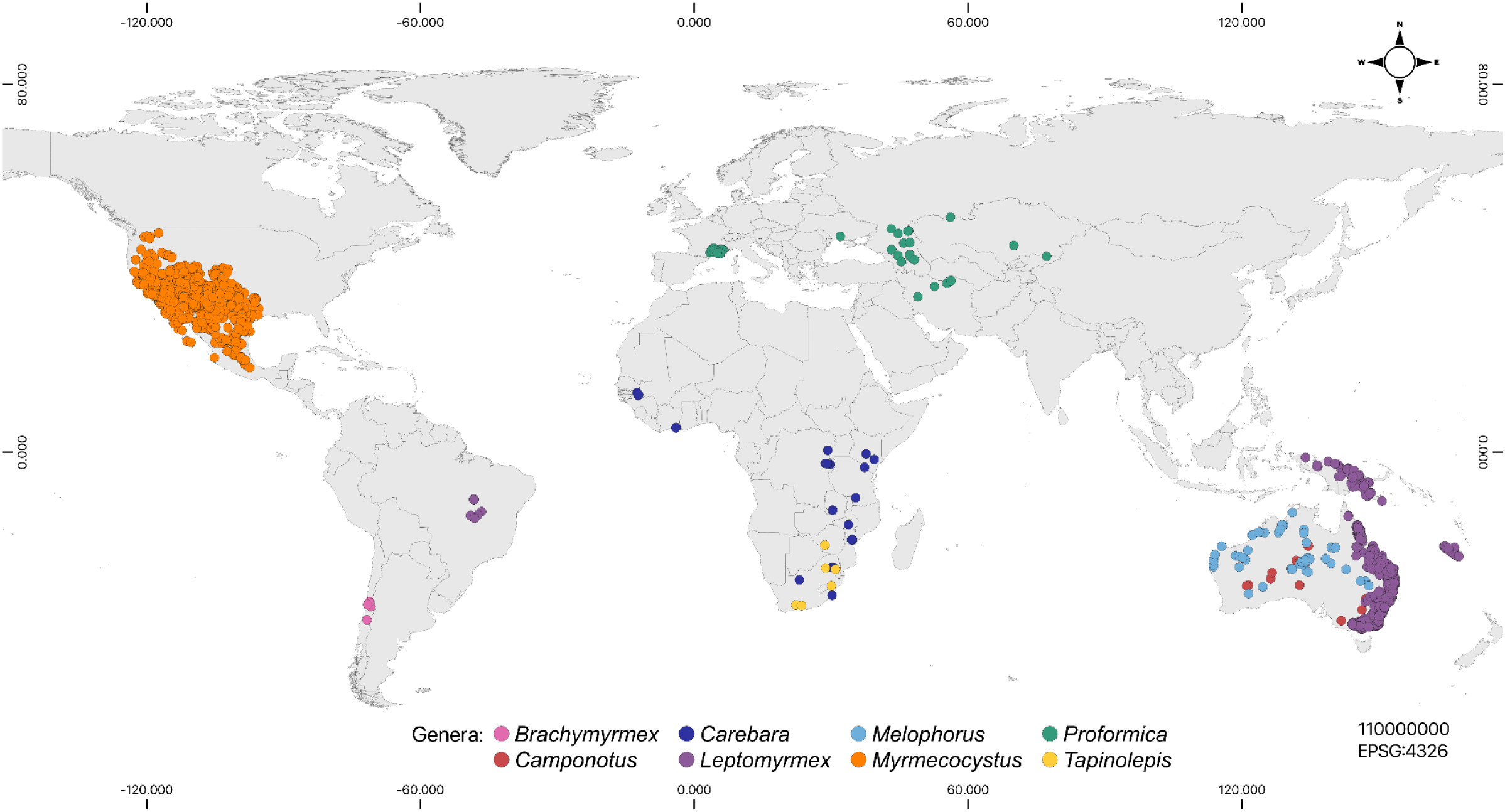
Global distribution of honeypot ants, highlighting the distribution of each genus. Plotted are cleaned occurrences used for all analysis. *Brachymyrmex*, *Camponotus*, *Carebara*, *Melophorus*, and *Tapinolepis* all represent single species. Occurrence data downloaded from GBIF (Derived dataset 2025), including personal collections.

### 3.2 Species ensemble models and predictors importance

Ensemble models showed good predictive performance overall (EMwmean: AUCprg = 0.965 ± 0.153, range = 0.856-1.000; TSS = 0.889 ± 0.085, range= 0.707-1.000; Figure S2). With one exception for *Myrmecocystus tenuinodis*, which had a negative ensemble AUCprg (−0.069), possibly due to its limited occurrence record (n = 8; Figure S3). This species was kept in subsequent analyses for completeness but its results are interpreted with caution. Individual models, which totalled 1708 across all species, showed a generally high calibration performance (mean AUCprg = 0.912 ± 0.282). Richness maps showed a high of 12 species co-occuring in Australasia and 16 in Nearctic (Figure 4), with higher richness closer to coastal areas and dry areas in North America.

**Figure 4.**
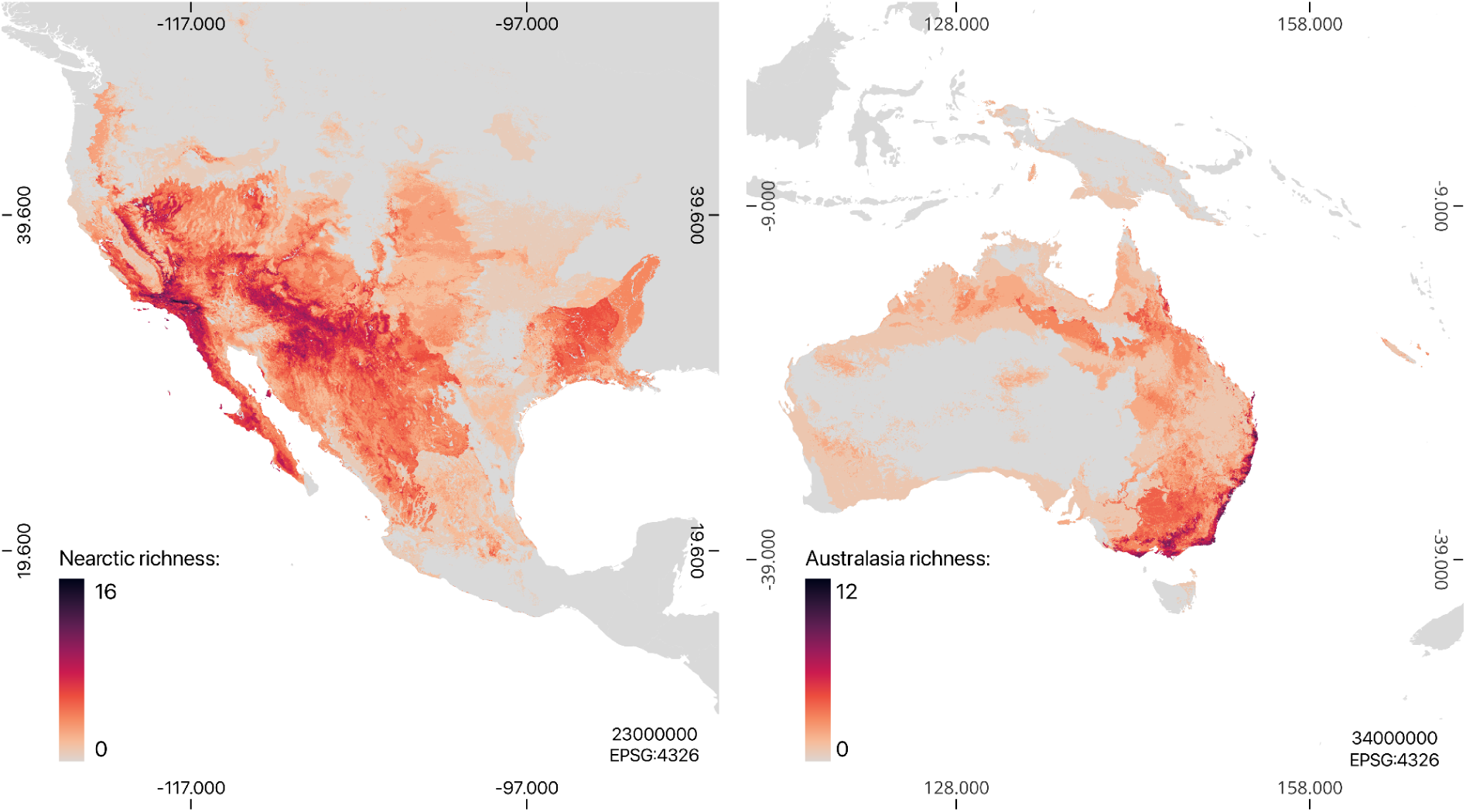
Predicted richness reached a maximum of 16 co-occurring species in the Nearctic (*Myrmecocystus*; left), and 12 in Australasia (*Leptomyrmex*, *Camponotus*, and *Melophorus*; right).

The three highest-ranked variables overall were precipitation of the driest quarter (bio19), precipitation of the coldest quarter (bio17), and mean diurnal soil temperature range (SBIO2; Figure 5). At the species level, tree cover and SBIO2 were the top predictor for six species each, followed by mean soil temperature of the wettest quarter (SBIO8) for five species (Figure 5).

**Figure 5.**
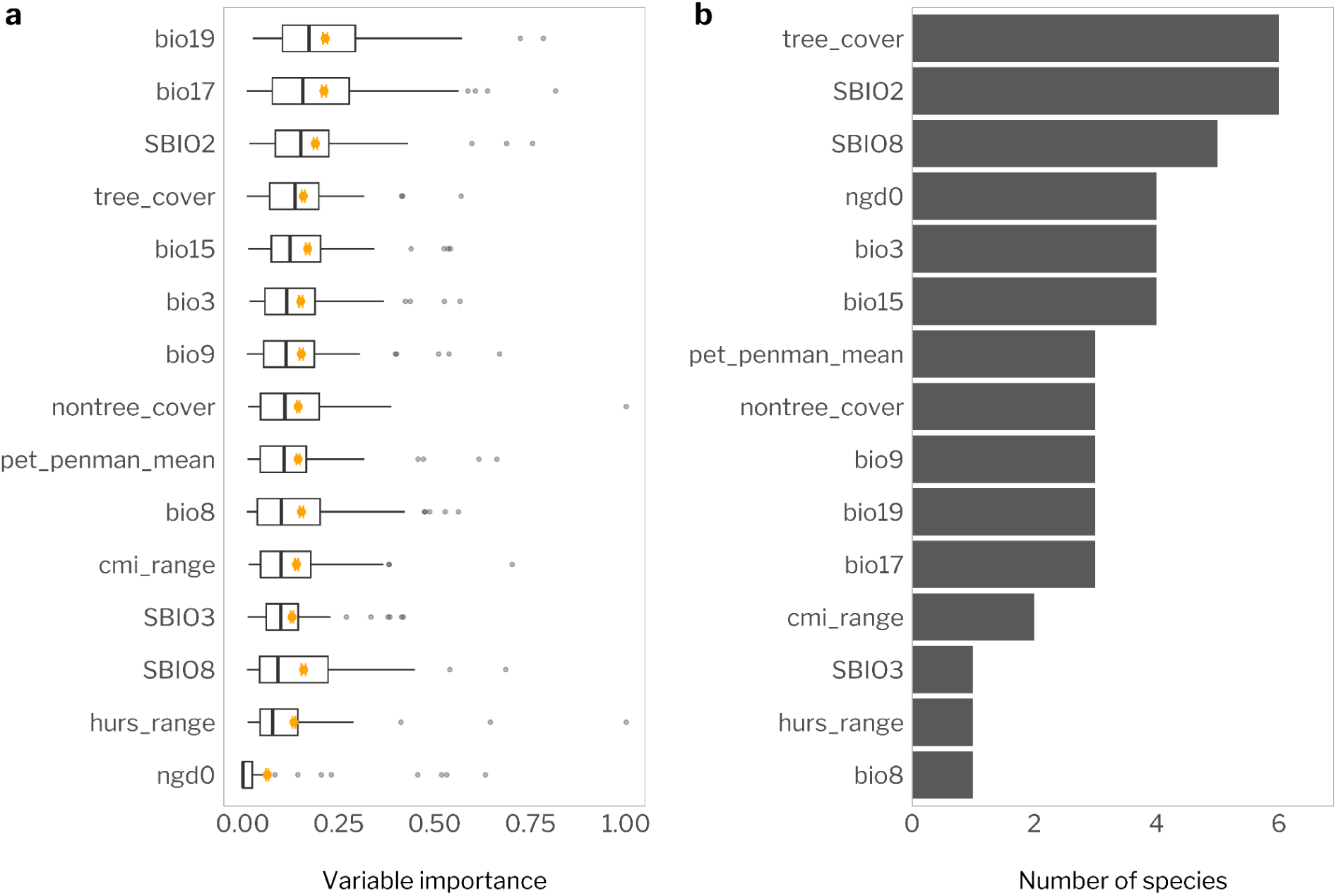
Precipitation of the coldest (bio17) and driest (bio19) quarters were the most important predictors overall (a). Of all variables, tree cover and SBIO2 were the most important variables for six species each (b). Boxplots show variation across ensemble models, with orange points indicating mean ± SD; and black bars show each time variables were considered the most important contributor across species.

Importance profiles revealed two broad clusters regardless of genus or region (Figure 6). One cluster was characterized by higher importance of growing degree days (ngd0), potential evapotranspiration (pet_penman), and SBIO2; the other by higher tree cover, SBIO8, and mean temperature of the driest quarter (bio9), with lower ngd0. Consistent with this pattern, neither genus nor region strongly structured importance profiles: genus was not significant (df = 7, R2 = 0.170, f = 1.205, p = 0.162), and region, while significant, explained little variation (df = 4, R2 = 0.138, f = 1.761, p = 0.019). Both groupings showed heterogeneous within-group dispersion (p < 0.01), and pairwise comparisons identified differences only between Australasia-Nearctic and Neotropic-Australasia pairs (p < 0.05; Table S3).

**Figure 6.**
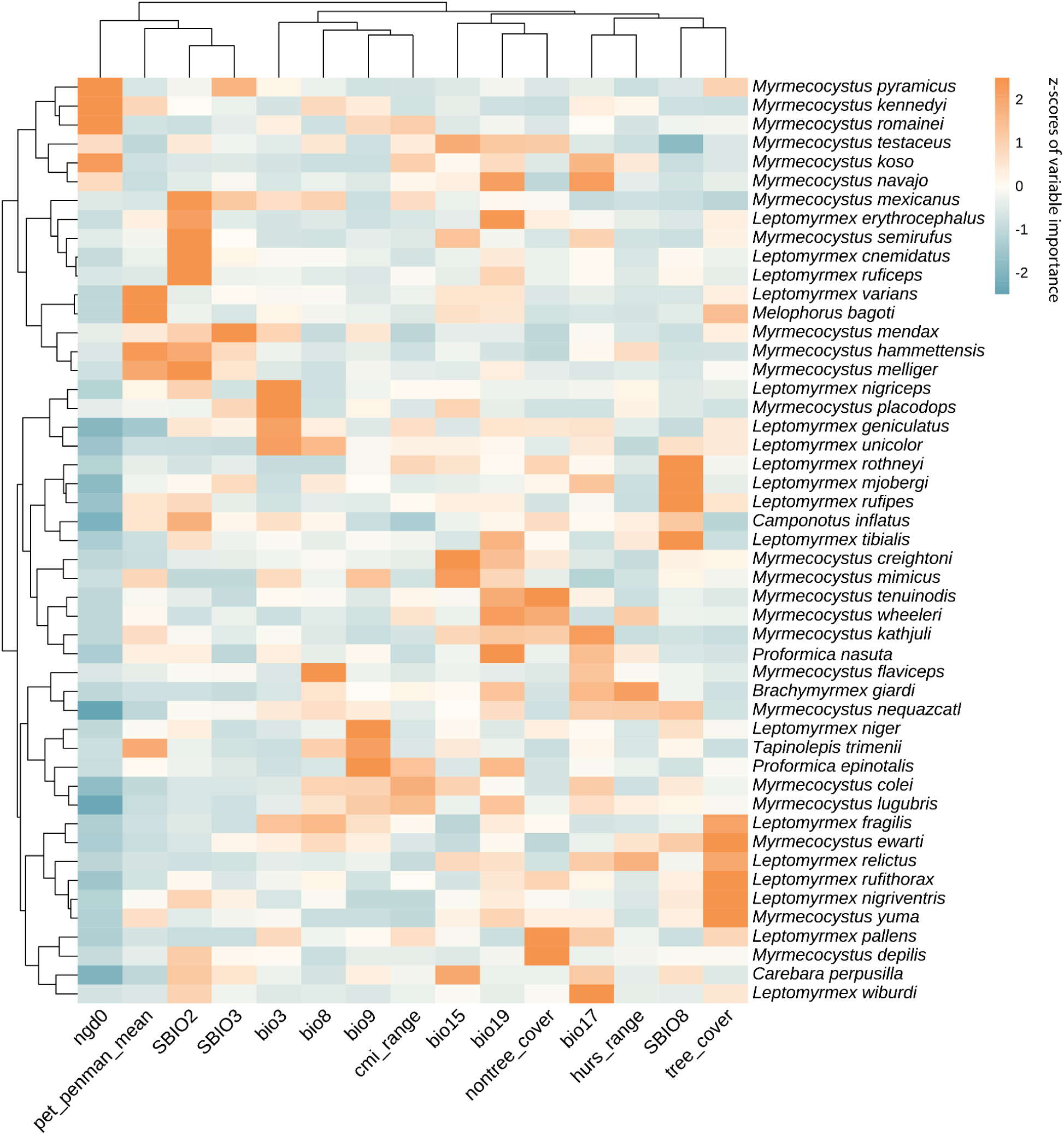
Species clustered into two major groups based on predictor importance, although no consistent genus-level pattern was observed. Heatmap shows variable importance values standardized within species (z-scores) before clustering to emphasize relative rather than absolute importance. Species and variables were clustered using Pearson correlation distance. High values indicate variables that are important relative to other predictors for a given species, not across all species globally.

For the Australasia-Nearctic comparison, importance profiles differed significantly only for ngd0, with higher importance in the Nearctic (D = 0.867, p < 0.001). To further explore whether differences in importance profiles reflect differences in actual environmental conditions, we computed Cohen’s d for environmental values extracted from occurrence points. Environmental values for ngd0 were nearly identical between regions despite the importance difference. Eight variables showed large-to-huge environmental differences, led by tree cover, SBIO2, bio17, and bio19 (Figure 7; Table S4). For the Neotropic-Australasia comparison, no importance profile differences were significant, yet four variables showed significant environmental differences: bio19, nontree_cover, bio15, and bio17 (Figure 7; Table S4).

**Figure 7.**
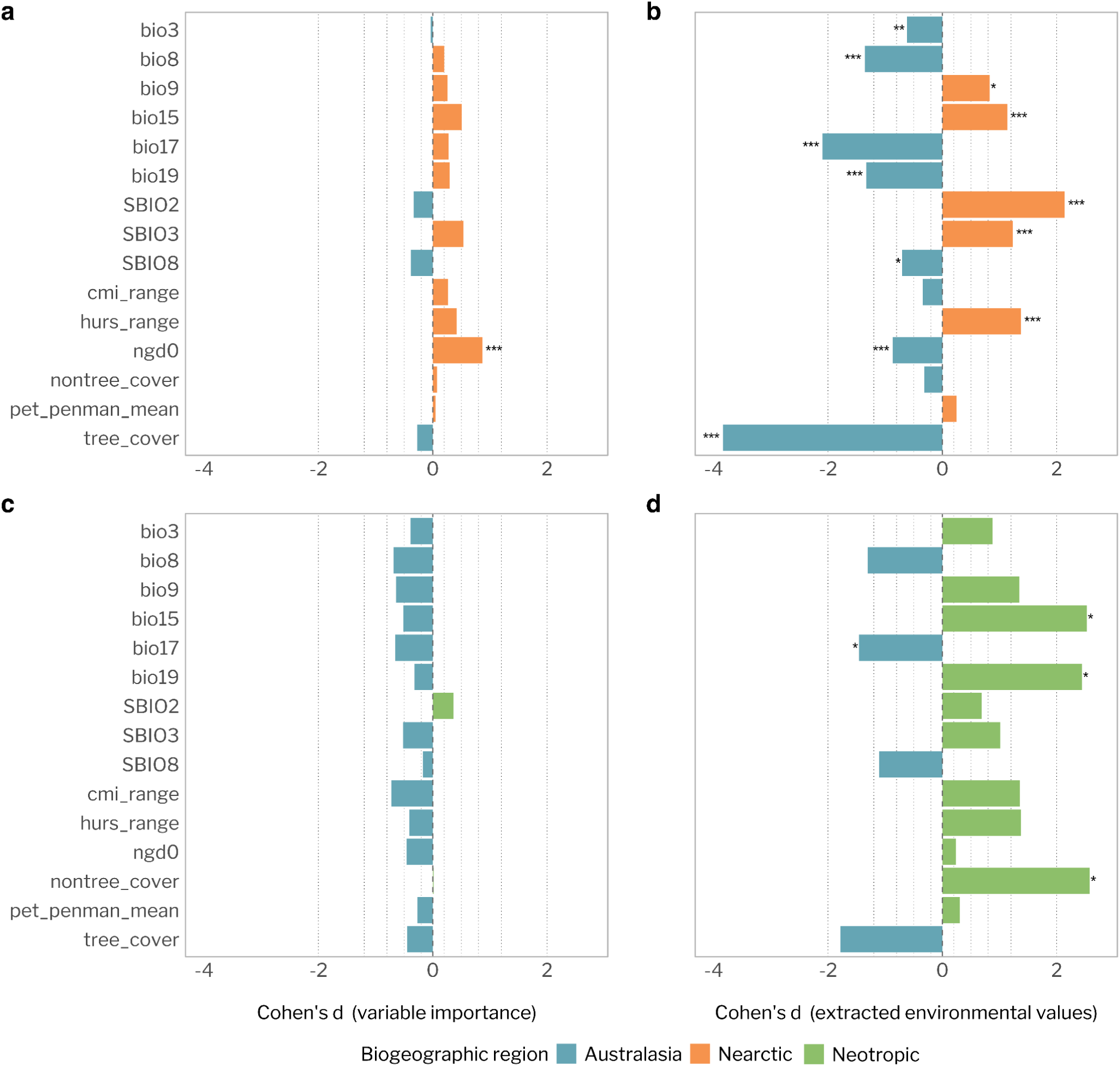
Australasian and Nearctic honeypot ants rely on similar predictors but occupy different environmental conditions. Cohen’s d effect sizes comparing variable importance profiles (a, c) and occurrence-derived environmental values (b, d) between regions. Bars are colored by the group with higher values; significance assessed by Wilcoxon rank-sum tests (*** p < 0.001; ** p < 0.01; * p < 0.05). Dashed lines indicate effect size thresholds (Sawilowsky 2009).

Variable importance profiles were not associated with geographic distance among species, either globally (Mantel r = 0.016, p = 0.299) or after controlling for biogeographic region (partial Mantel r = −0.035, p = 0.819). As the Nearctic fauna is dominated by *Myrmecocystus* and Australasia by *Leptomyrmex*, these genera account for most of the among-species variation in importance profiles.

### 3.3 Multivariate niche analysis

#### 3.3.1 Species environmental space

Comparing Boolean pixels against background conditions, 47 species showed significant differences in centroid position and 41 in dispersion (PERMANOVA and PERMADISP: median p < 0.05; Table S5; Figure S4). Species without significant differences were among those with fewest records (*Myrmecocystus colei* Snelling, 1976 and *M. kathjuli* Snelling, 1976, n = 8). When occurrence records were compared against the background, fewer species showed significant centroid differences (n = 31) or dispersion differences (n = 16), likely reflecting reduced statistical power from smaller sample sizes.

Despite these differences relative to background, occurrences and Boolean pixels were largely consistent with each other: 29 species showed no significant centroid differences and 37 showed no significant dispersion differences when compared directly (PERMANOVA and PERMADISP: median p > 0.05). Where significant differences occurred, effect sizes were small (R² < 0.1), and visual inspection confirmed broad overlap in PCA space.

#### 3.3.2 Within biogeographic regions comparisons

All within-region comparisons with occurrence data showed significant differences in species centroid positions in environmental space (PERMANOVA: median p < 0.05; Table S6), with species identity explaining considerable variation in most regions (Table S6). Likewise, within-group dispersion was significant in all regions (PERMDISP: median p < 0.05; Table S6). We observed a similar pattern using both Boolean suitable pixels and occurrence points (Table S6).

##### 3.3.2.1 Australasia

In the Australasian region, the first two PCA axes explained 58.4% of environmental variation (PC1 = 35.9%, PC2 = 22.6%). PC1 contrasted open, high-evapotranspiration environments with greater diurnal temperature variation (positive: SBIO2, nontree_cover, pet_penman, SBIO8) against wetter, more vegetated conditions (negative: bio19, bio17, tree cover, bio3). PC2 reflected seasonal climatic variation, with negative values associated with precipitation seasonality, temperature of the driest quarter, and moisture variability (bio9, pet_penmean, bio15, cmi_range, and SBIO8).

Species were clearly structured along these axes. *Camponotus inflatus* and *Melophorus bagoti* (Figure 8) occupied negative PC1 values, consistent with open, thermally variable environments, while most *Leptomyrmex* species clustered toward central to positive PC1 values, reflecting wetter and more tree-covered conditions. Within *Leptomyrmex*, most species overlapped broadly, though *L*. *niger* and *L*. *fragilis* were positioned toward high positive values on both axes, indicating occupancy of wetter, more seasonally variable environments.

**Figure 8.**
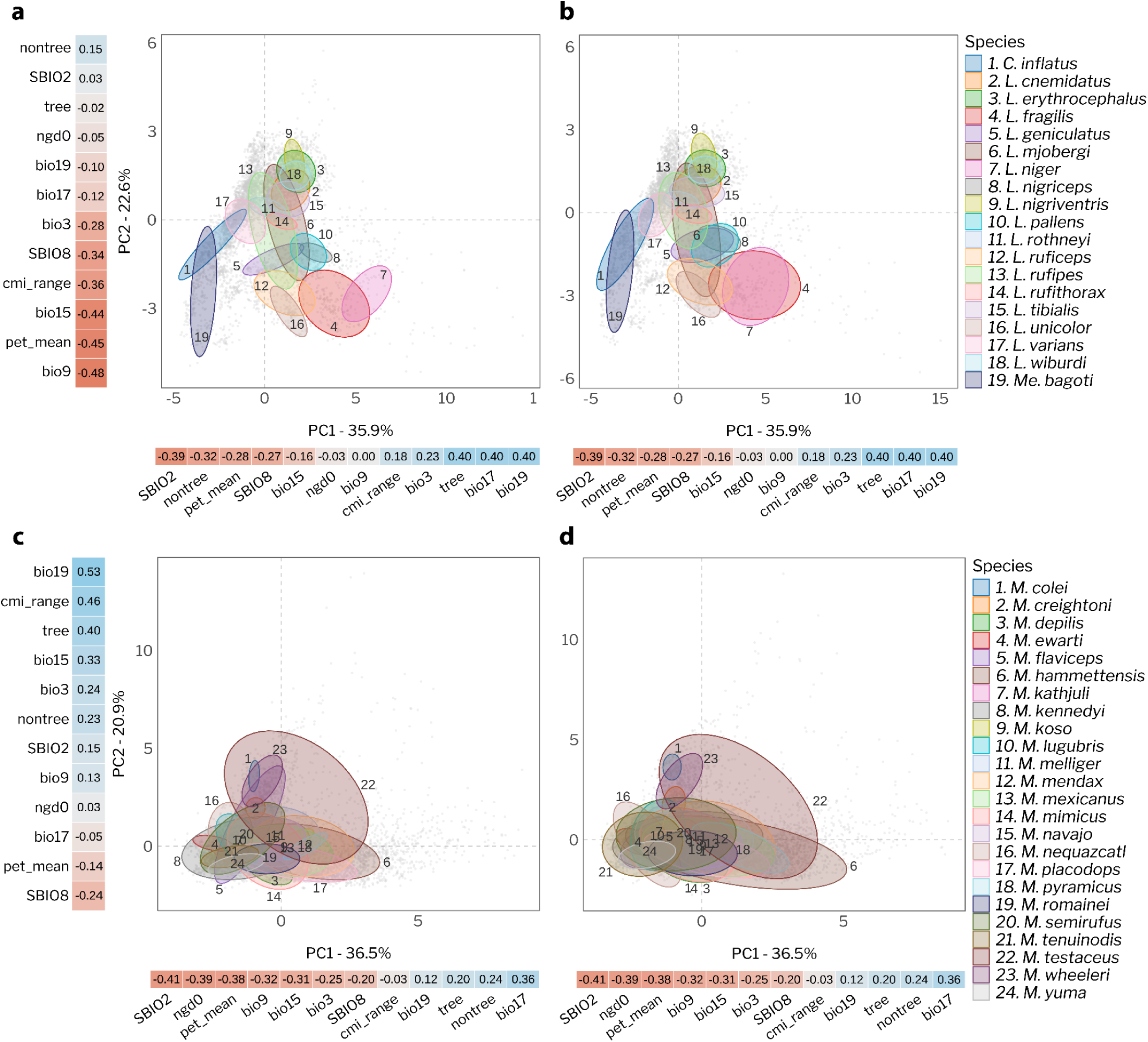
Environmental PCA spaces for Australasian (a,b) and Nearctic (c, d) honeypot ants. Boolean suitable pixels (a, c) and occurrence records (b, d) projected into the first two environmental PCA axes. Gray points represent background environmental conditions within the accessible area, while colored 75% confidence ellipses represent species environmental spaces. Color bars indicate PC loadings for each predictor variable and for each PC axis; blue represents positive values and red negative values.

Most pairwise PERMANOVA comparisons were significant (97.7% of 171 pairs, p < 0.05; Table S7), indicating widespread environmental differentiation across *Leptomyrmex*. Non-significant pairs corresponded to the highest overlap values (Figure 10), with species pairs such as *L. fragilis*-*L. niger*, *L. cnemidatu*s-*L. tibialis*, and *L. geniculatus*-*L. pallen*s showing high Schoener’s D and low centroid distances. *L. erythrocephalu*s and *L. wiburd*i were an exception: despite a significant PERMANOVA (R² = 0.011), they showed high overlap (D = 0.74) and low centroid distance, suggesting their separation reflects niche breadth differences rather than distinct environmental positions. *L. rufithorax* showed the broadest range along PC1, while species in New Guinea and New Caledonia occupied overlapping positions along PC2.

Geographic distance was a strong predictor of environmental differentiation in Australasia (GAM: adj. R² = 0.574, p < 0.001), with geographically distant species tending to occupy more distinct environmental spaces.

##### 3.3.2.2 Neartic

For the Neartic region, the first two PCA axes explained 57.4% of environmental variation, with the first seven axes retained for multivariate analyses (91.2%). PC1 contrasted open, thermally variable, high-evapotranspiration environments (negative: SBIO2, pet_penman, ngd0, bio9, bio15, SBIO8) against wetter, more vegetated conditions (positive: bio17, bio19, tree_cover, nontree_cover). PC2 reflected moisture and seasonal variation, with positive values associated with bio19, cmi_range, tree_cover, bio15, and bio3, and negative values with SBIO8 and pet_penman.

Most *Myrmecocystus* species pairs were significantly differentiated in environmental space (90.2% of 276 pairs, mean R² = 0.17; Table S7). Non-significant pairs showed consistently higher overlap and lower environmental centroid distances than significant pairs (median Schoener’s D: 0.44 vs. 0.30; median centroid distance: 0.98 vs. 2.48). Although non-significant pairs tended to have smaller sample sizes, the pattern held after accounting for this, suggesting that some significant comparisons reflect both real differentiation and greater statistical power from larger samples.

As in Australasia, geographic distance was positively associated with environmental differentiation (GAM: adj. R² = 0.241, p < 0.001), though the relationship was weaker and less linear, suggesting finer-scale ecological partitioning among Nearctic species.

##### 3.3.2.3 Palearctic, Afrotropic and Neotropic

In the three remaining regions, each containing only two species, all pairs showed strong environmental differentiation (PERMANOVA: p < 0.01, R² = 0.45-0.49; Schoener’s D < 0.01 for all). PCA axes explained 63.4-68.8% of variation in the first two components, with six axes retained for multivariate analyses in each region (Figure 9). Across all three regions, PC1 consistently separated species along a gradient from open, thermally variable, and seasonally dry environments toward wetter and more vegetated conditions, while PC2 reflected climatic seasonality and moisture variability.

**Figure 9.**
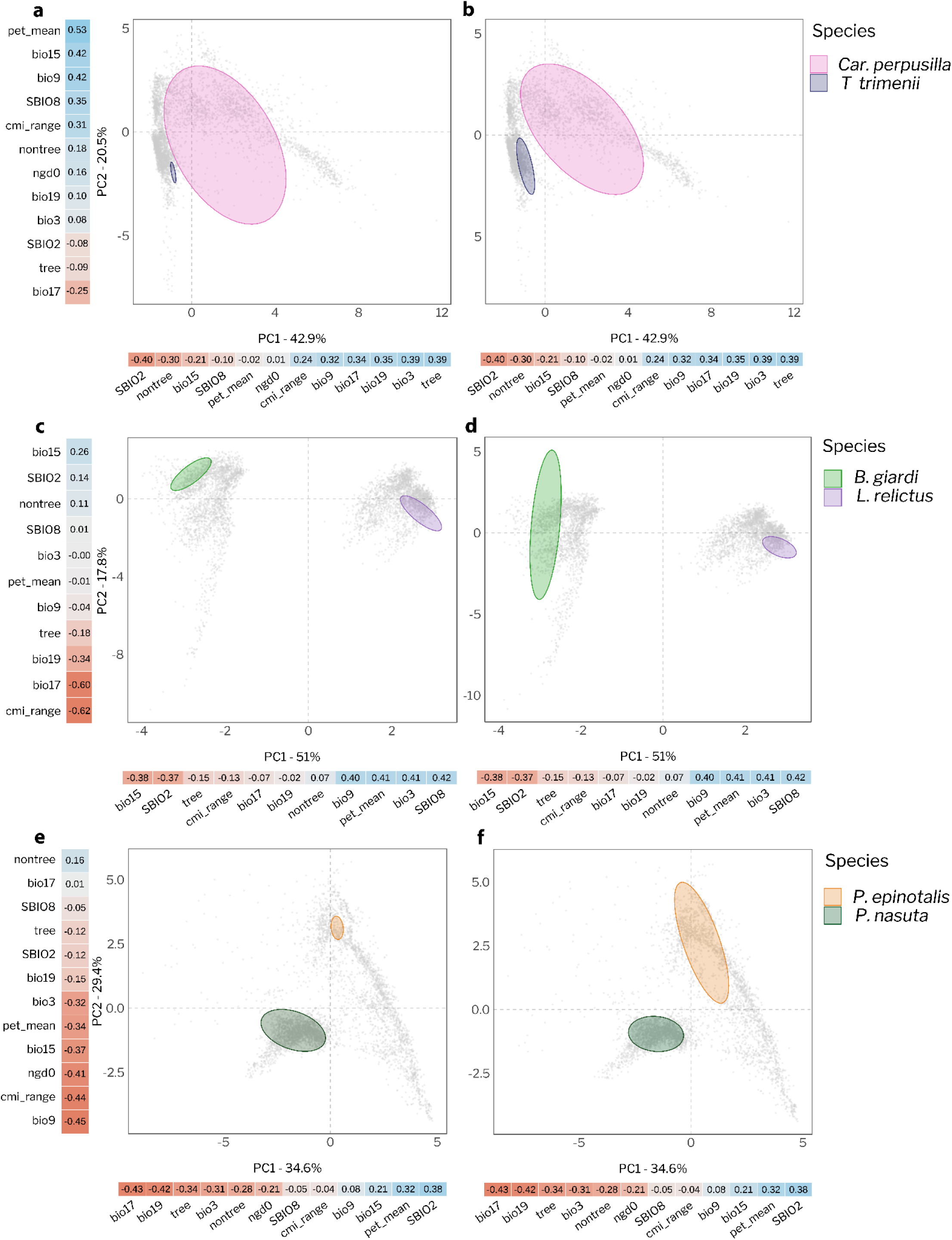
Environmental PCA spaces for Afrotropic (a,b), Nearctic (c,d), and Palearctic (e,f) honeypot ants. Boolean pixels (a, c, e) and occurrence records (b, d, f) projected into the first two environmental PCA axes. Gray points represent background environmental conditions within the accessible area, while colored 75% confidence ellipses represent species environmental spaces. Color bars indicate PC loadings for each predictor variable and for each PC axis; blue represents positive values and red negative values.

Within each region, species occupied opposite ends of this gradient (Figure 9). In the Palearctic, *P. epinotalis* was associated with greater diurnal temperature variation and higher evapotranspiration, while *P. nasuta* occupied wetter, more vegetated environments with stronger moisture variability and frost-related conditions. In the Afrotropic, *T. trimenii* was positioned toward open, thermally variable conditions with stronger seasonality, while *C. perpusilla* occupied broader, wetter, and more vegetated environmental space. In the Neotropic, *B. giardi* was associated with seasonally variable, moisture-structured environments, while *L. relictus* occupied warmer, more energy-demanding conditions. In all cases, species were also geographically separated, suggesting that geographic distance contributes to their environmental distinctiveness.

#### 3.3.3 Between biogeographic regions comparisons

Pairwise PERMANOVA on species environmental centroids revealed significant differences between Neotropic-Australasia, Neotropic-Nearctic, Australasia-Nearctic, Afrotropic-Nearctic, and Nearctic-Palearctic (occurrence data, BH-adjusted p < 0.05; Table S8). Results from Boolean data were broadly consistent, though Neotropic-Australasia and Afrotropic-Nearctic were not significant.

Environmental centroid distance was positively associated with geographic distance (Mantel r = 0.44, p = 0.001; MRM: R² = 0.215, p < 0.0001), with a nonlinear relationship (GAM: adj. R² = 0.495, edf = 8.8, p < 0.0001). However, when region membership was included, geographic distance lost its effect and only regions membership remained significant (MRM: R^2^= 0.244; geographic distance: p = 0.50; region: β = 2.08, p = 0.002), indicating that the apparent distance-decay pattern is driven by broad among-region differences rather than a continuous geographic effect.

In the global PCA (PC1-PC2: 51.7% of variation; Figure 10), Nearctic species clustered in positive PC1 space, reflecting association with open, thermally variable, high-evapotranspiration environments (SBIO2, pet_penman, bio15, bio9), while Australasian, Neotropical, and Palearctic species occupied negative PC1 space, associated with wetter and more vegetated conditions (bio19, tree_cover, bio3, cmi_range). Most Afrotropical species followed a similar pattern, with the exception of *Tapinolepis trimenii*, which clustered alongside Nearctic species in positive PC1 space, as did three Australasian species (*Camponotus inflatus*, *Melophorus bagoti,* and *Leptomyrmex varians*).

**Figure 10.**
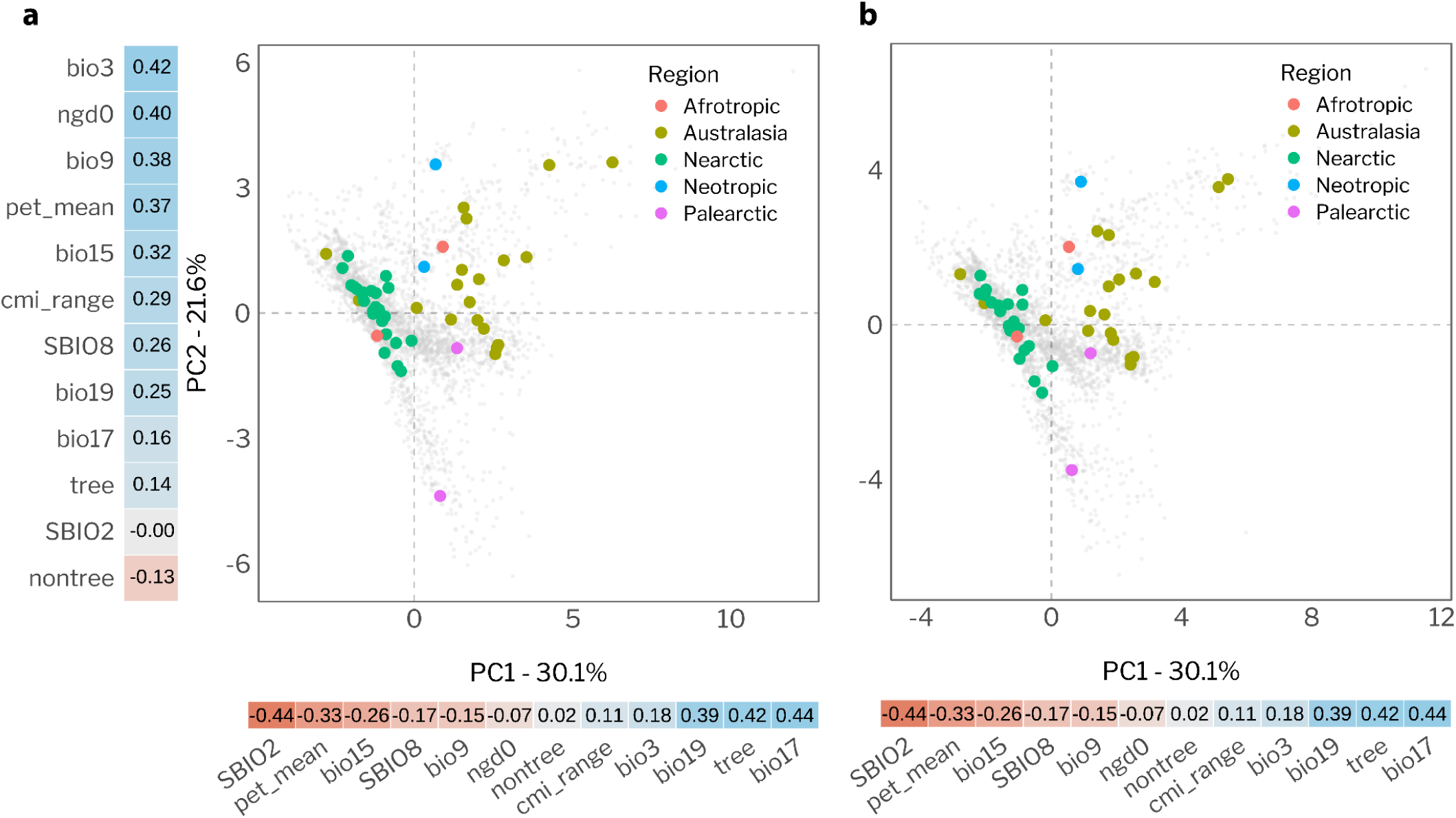
PCA of mean climatic space occupied by honeypot ant species across biogeographic regions for Boolean (a) and occurrence (b) data. Gray points show the full background environmental space.

Of 726 between-region pairwise comparisons, 724 were significant (PERMANOVA p < 0.05), with generally very low overlap (median Schoener’s D = 0.045). The two non-significant pairs were *Camponotus inflatus*-*Myrmecocystus nequazcati* (D = 0.18) and *Tapinolepis trimenii*-*M. melliger* (D = 0.54; Figure 11), the latter indicating substantial shared environmental space despite a shift in multivariate position. Beyond these cases, high overlap values (D ≥ 0.5) occurred in several between-region pairs outside Australasia-Nearctic, with the highest being *T. trimenii*-*M. romainei* (D = 0.65; Figure 11), suggesting partial environmental convergence across regions. Results from Boolean data were highly consistent with occurrence-based analyses throughout (Table S8).

**Figure 11.**
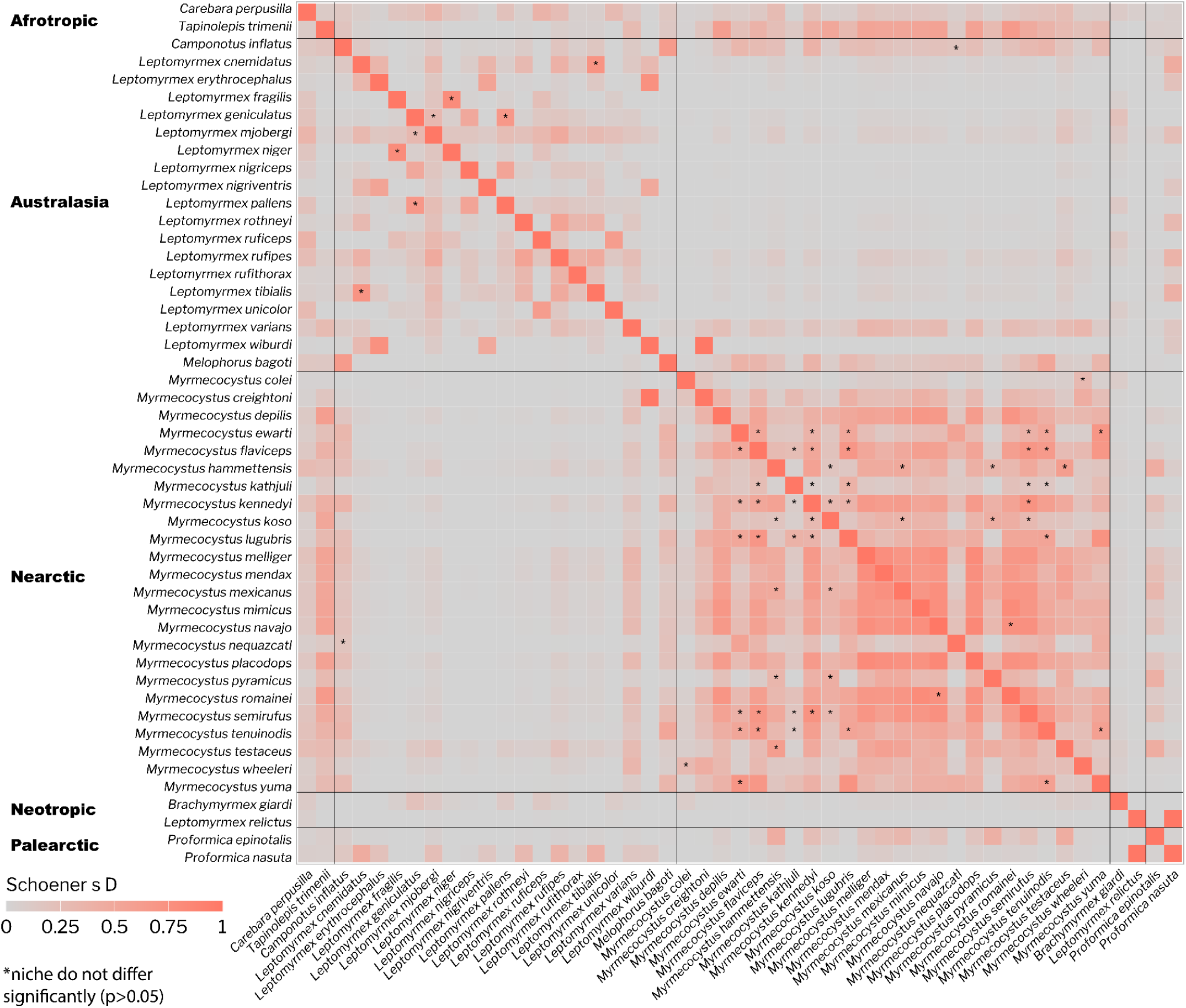
Niche overlap is high, especially between Afrotropic-Nearctic and within regions. Heatmap showing Schoener’s D values for species pairwise comparisons with occurrence data. * indicates pairs that PERMANOVA was not significantly different. Black lines separate species by biogeographical region.

## 4 DISCUSSION

Repletism is a convergent morphological trait (Nogueira et al., 2026) however, contrary to our initial hypothesis, the current climatic and environmental conditions are not shared among all honeypots as a group. Our ensemble models showed high overall predictive performance, indicating that the selected variables effectively characterize honeypot ant environmental niches. The broad overlap between occurrences and Boolean pixels suggests that binary predictions captured a restricted but representative subset of available environmental conditions. Using Boolean points proved to be appropriate for niche comparisons without treating them as exact representations of species distributions, and enhancing the power of our results.

### 4.1 There is no pattern in variable importance profiles among the honeypots

Overall, variable-importance profiles showed limited taxonomic or predictor structure. This indicates that no consistent set of environmental variables was associated with honeypot ants across genera or biogeographic regions (Muñoz et al., 2016; Smith & Santos, 2020). Our results show that predictors, especially precipitation and water-availability related, can contribute similarly to species distributions while species occupy different conditions along those gradients. Variable-importance profiles are not direct measures of environmental similarity, but we show here that they can serve as informative ENM construction summaries for identifying the environmental dimensions most relevant to multivariate niche analyses.

### 4.2 Honeypot ant niche differences within regions

Within regions, species consistently separated along a gradient from open, thermally demanding conditions to wetter and more vegetated environments. This pattern reflects both local climatic gradients and lineage-specific ecology. In Australasia, the strong geographic-environmental distance relationship suggests that species distributions follow the continent’s broad climatic structure, from arid interiors to tropical and temperate margins (Andersen, 1995); and this is also consistent with honeypots’ known ecologies (Christian & Morton, 1992; Lucky & Ward, 2010). Notably, the highest niche overlap with non-significant pairwise differences occurred among coastal and island species in New Guinea and New Caledonia. This likely reflects climatically homogeneous conditions and limited dispersal inherent to island systems, which constrain environmental differentiation among co-occurring taxa (Losos & Ricklefs, 2009; MacArthur & Wilson, 2001).

In the Nearctic, despite all species belonging to a single arid-adapted genus, *Myrmecocystus* showed finer-scale partitioning along gradients of precipitation, vegetation cover, and moisture availability. This is consistent with the weaker geographic-environmental distance relationship in the Nearctic relative to Australasia. This suggests that local ecological factors matter more than geographic distance alone. Competition is one explanation, as closely related species are expected to diverge in habitat use to avoid exclusion (Pigot & Tobias, 2013; Sfenthourakis et al., 2006). Physiological constraints likely reinforce this pattern, as ants forage close to their thermal limits (Cerdá et al., 1998; Lighton & Turner, 2004; Roeder et al., 2021), and small differences in temperature and moisture can translate into meaningful ecological boundaries, particularly where water balance is a key constraint (Bujan et al., 2016; Parr & Bishop, 2022).

In the remaining regions, each represented by only two species, a similar gradient consistently separated species. Together, these patterns show that honeypot ants occupy markedly different environments even within the same region, setting the stage for understanding how these differences scale across realms.

### 4.3 Honeypot ant niche differences across regions

The clear environmental structure recovered across honeypot ant species globally indicates that repletism is associated with a wide range of ecological contexts, rather than a single shared environment. The separation of open, thermally demanding habitats from wetter, more vegetated conditions is consistent with macroclimatic patterns across the regions and biomes where honeypot ants occur (Christian & Morton, 1992; Fernández-Escudero & Tinaut, 1998; Fischer et al., 2015).

The strongest between-region contrast was between the Nearctic and Australasia. Despite this, both showed comparable values for potential evapotranspiration and growing degree days (ngd0). Ngd0, which reflects accumulated thermal energy, drives phenological processes including plant growth and insect activity (McMaster & Wilhelm, 1997). In sparsely vegetated Nearctic habitats, ngd0 may indirectly capture seasonal nectar availability, a primary resource stored by Myrmecocystus repletes (Conway, 1986; Meurville et al., 2025). Australasian ants were associated with high evapotranspiration, which reflects thermally demanding atmospheric conditions even in the absence of strict aridity. For ground-nesting ants closely tied to soil microclimate, such conditions may favor fluid storage regardless of drought (Bujan et al., 2016; Cerdá et al., 1998; Lighton & Turner, 2004). Given that repletes may function in water storage as well as carbohydrate storage (Conway, 1977, 1986), shared thermal demand across climatically distinct regions may help explain the occurrence of repletism outside strictly arid environments.

Interestingly, *Leptomyrmex relictus*, the only extant *Leptomyrmex* outside Australasia, occupied a distinct environmental space and showed little overlap with other *Leptomyrmex*. This suggests that niche conservatism does not fully explain *Leptomyrmex* distributions (Wiens et al., 2010), and that some lineages may have retained repletism while colonizing different environments (Boudinot et al., 2016; Sawh et al., 2023). *Tapinolepis trimenii* occupied environmental space strongly overlapping with Nearctic *Myrmecocystus* despite being geographically and phylogenetically distant, suggesting that desert-like conditions can independently produce similar environmental associations across lineages.

### 4.4 Repletism may solve different ecological problems across lineages

Convergence is often associated with similar environmental pressures, but similar traits can also evolve under distinct selective contexts when they solve different ecological or physiological challenges across lineages (Losos, 2011). Succulent plants, for instance, exhibit convergent morphologies despite occupying climatically distinct regions, a pattern attributed to shared functional demands rather than shared environments (Alvarado-Cárdenas et al., 2013). Our results suggest that honeypot ants may represent a similar case.

The patterns we observed might also reflect lineage-specific differences in behavior, physiology, morphology, and colony organization (Jaekel & Wake, 2007; Morris et al., 2019; Vidal-García & Keogh, 2015). While life history details remain sparse for most genera in this study, we know that *Leptomyrmex* repletes are less specialized and more mobile than those of *Myrmecocystus* (Nogueira et al., 2026; Plowman, 1981). That could mean that repletes might have slightly different functions and demands in different species, driven by different environmental pressures. *Leptomyrmex* is also the only honeypot lineage within Dolichoderinae, a subfamily with distinct colony-level traits that may independently shape environmental associations (Arnan et al., 2012; Ward et al., 2010). Together, these results suggest that repletism may have evolved and persisted under different combinations of environmental conditions across lineages, shaped by lineage-specific ecology, resource use, physiology, and historical contingency.

### 4.5 Future directions

Our results show that contemporary environmental conditions do not provide a unifying explanation for the convergent evolution of repletism, pointing to the need for broader historical and biological perspectives. Incorporating past climate conditions and species biogeography will be our next step, as paleoclimatic reconstructions can shed light on the original environments and selective pressures under which repletes first evolved (Ralph & Coop, 2015; Rosenblum, 2006; Sakamoto & Ruta, 2012). Fossil evidence and ancestral state reconstruction suggest repletism originated around 45 million years ago in the Eocene with *Leptomyrmex* (Sawh et al., 2022). Dating the emergence of repletism in other lineages would allow us to test whether independent origins match with global aridification events. Studying honeypot ants in relation to sister-species environmental space might also help us understand if honeypots as a group are under distinct pressure.

Understanding whether similar genetic and developmental pathways underlie replete formation across lineages will also be key, as shared molecular mechanisms would strengthen the case for deep functional convergence (Holekamp et al., 2013; Molet et al., 2012). Complementary histological and physiological studies of nutrient storage strategies and colony-level dynamics are also needed to characterize functional variation across species (Casadei-Ferreira et al., 2020; Zharkov & Dubovikoff, 2023). Finally, biotic interactions such as microbial symbiosis may impose additional selective pressures on replete evolution and deserve further investigation (Coleine et al., 2024; McCutcheon et al., 2009; Nguyen et al., 2026).

## 5 CONCLUSIONS

This study provides the first broad-scale analysis of environmental niche across honeypot ant lineages, revealing that honeypots are distributed in different climatic contexts. While arid habitats are strongly associated with repletism, honeypots have diversified into distinct environmental spaces. The lack of a shared contemporary niche across the honeypots challenges the assumption that repletism resulted from convergent selective environments, and points toward resource variability as a possibility. However, current distributions do not necessarily reflect the environments in which the ants first evolved repletism. To understand the true selective pressures that have led to repletism, past climate must be taken into account. By establishing the environmental context of extant honeypot ant diversity, this work lays the groundwork for future comparative studies of convergent evolution across independent lineages.

## Acknowledgements

We thank Dr. Angelo Soto-Centeno for help with the initial conceptualization of the project. Henrique Morais Menezes for field assistance and the collection of *Leptomyrmex relictus* in Brazil. This work was supported by NSF BRC-BIO grant 2312984 to LK.

## Author Contributions

Bianca R. Nogueira and Lily Khadempour conceived the ideas. Bianca R. Nogueira collected the data and, together with Omar D. Léon-Alvarado, designed the methodology and analyzed the data. Bianca R. Nogueira led the writing of the manuscript, but all authors contributed critically to the drafts and final version.

Statement on inclusion

This work brings together authors from diverse backgrounds, including two with South American and one with Middle Eastern origins, and was led and advised by women. The majority of the data was publicly available. Some collections were completed in Brazil, in the first author’s home country, in collaboration with a local researcher. Our global analysis highlights under-sampled regions and the need for stronger research partnerships.

## Data availability statement

Scripts are available at https://anonymous.4open.science/r/honeypots_niche_modeling-25BF/. Occurrence data were sourced from two GBIF exports: a filtered derived dataset (GBIF.org, 2026; https://doi.org/10.5281/zenodo.20056097, retrieved 28 February 2025) and a direct occurrence download (GBIF.org, 19 April 2026; https://doi.org/10.15468/dl.87wgac).

