## Supplementary material for "Convergently-evolved honeypot ants show mixed signs of niche divergence": FiguresS1-6_Supp.docx

### [**Supplementary material**](https://authors.wiley.com/author-resources/Journal-Authors/Prepare/manuscript-preparation-guidelines.html/supporting-information.html)

### **Title: Convergently-evolved honeypot ants show mixed signs of niche divergence**

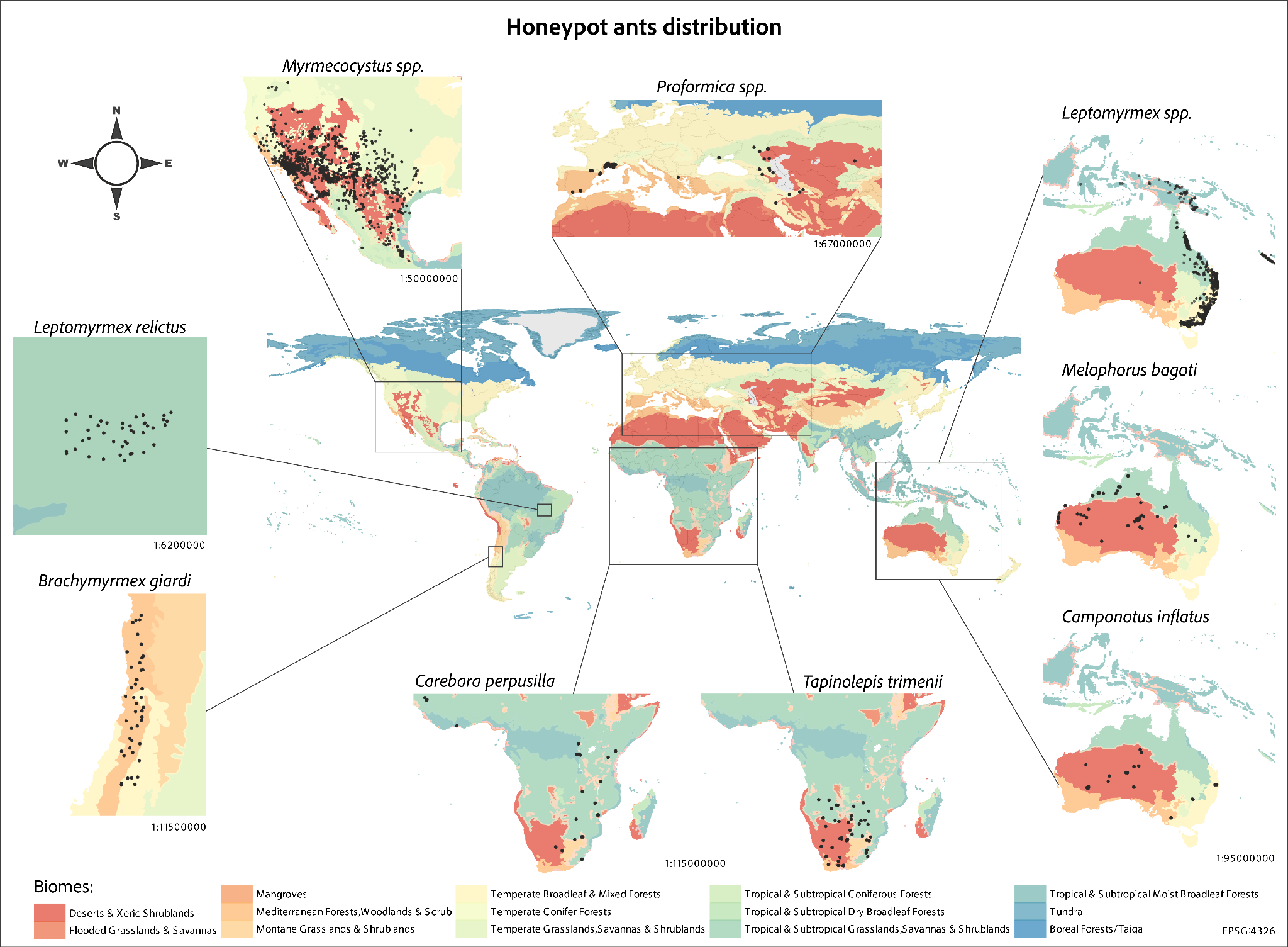

Figure S1. Global distribution of honeypot ants, highlighting the distribution of each genus and their occurrence across biomes (Dinerstein et al. 2017). For genera with multiple species, the points represent the combined occurrences of all species. Occurrence data downloaded from GBIF.

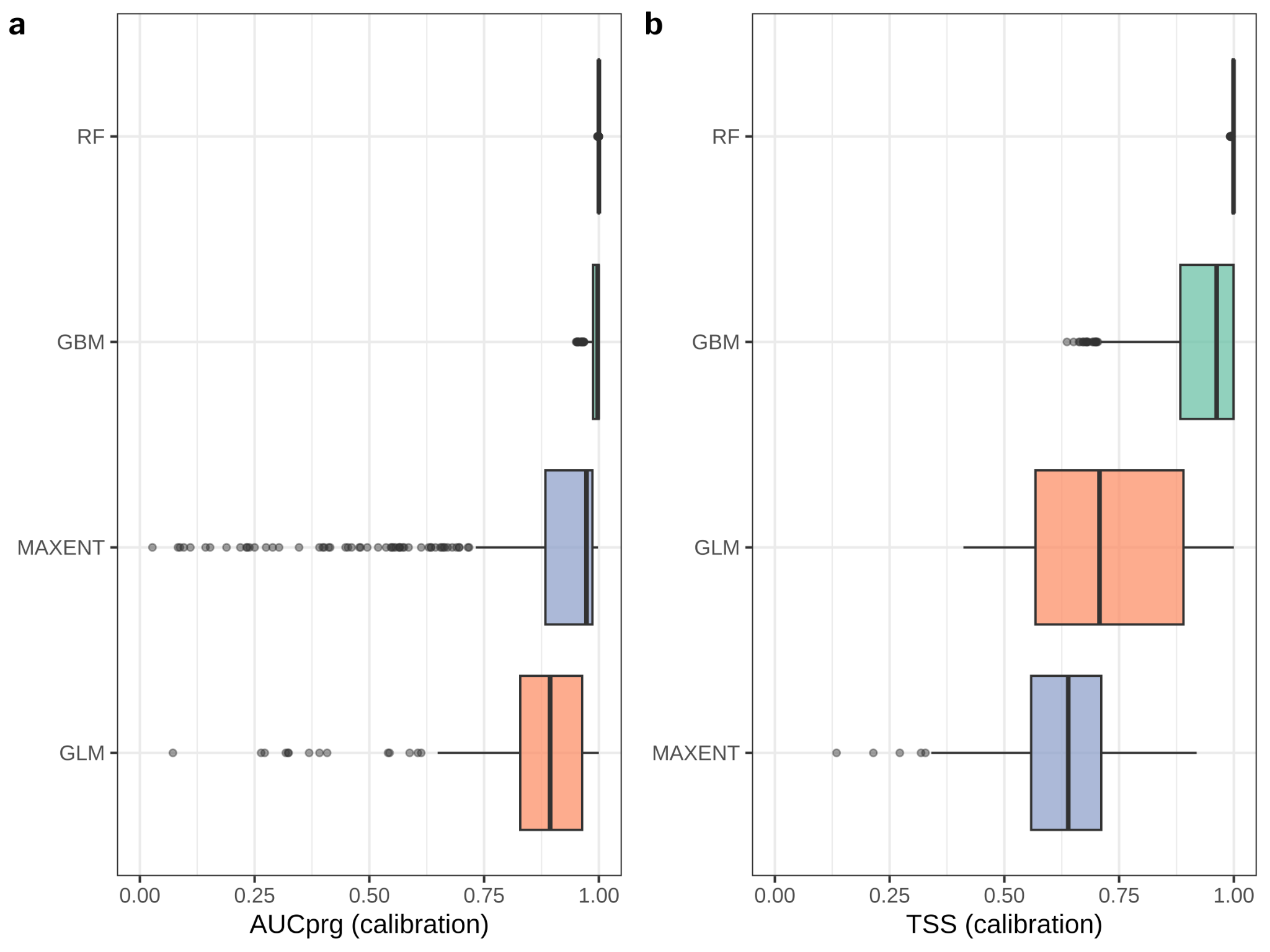

Figure S2. Individual models' calibration score by algorithm.

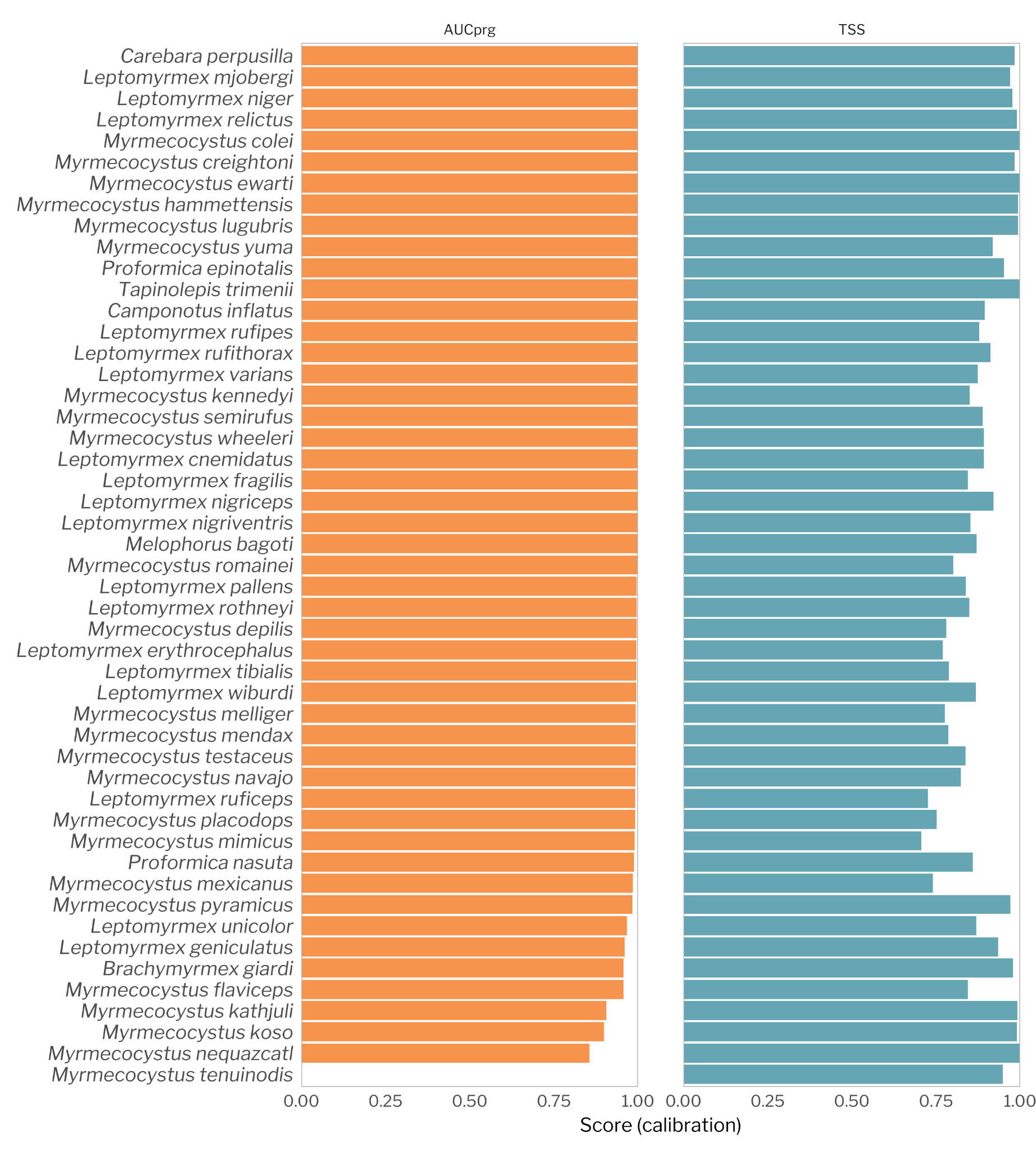

Figure S3. Ensemble models’ calibration score by species.

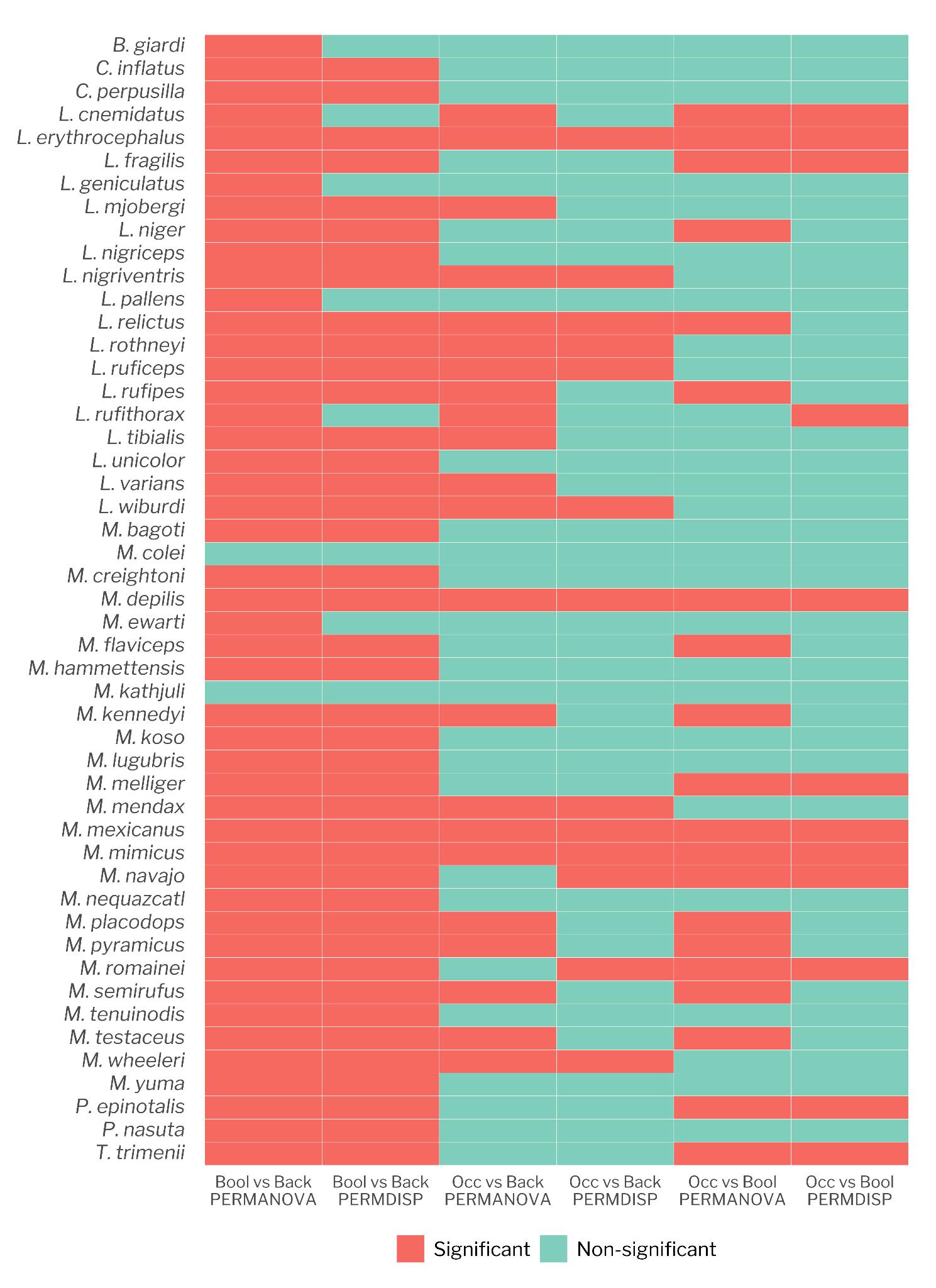

Figure S4. Heatmap to visualize the results between occurrence-boolean-background data.

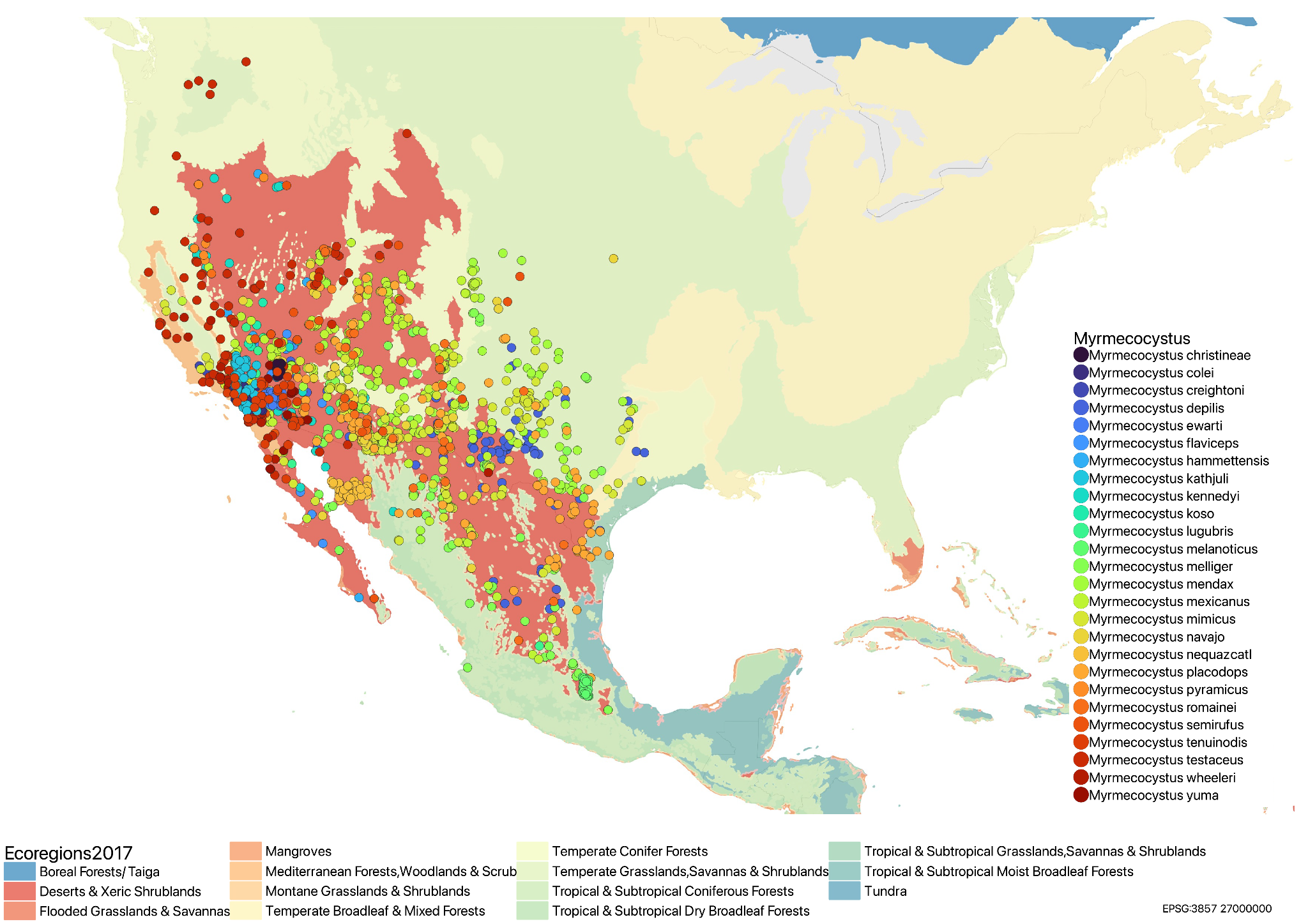

Figure S5. *Myrmecocystus* species occurrence points downloaded from GBIF.

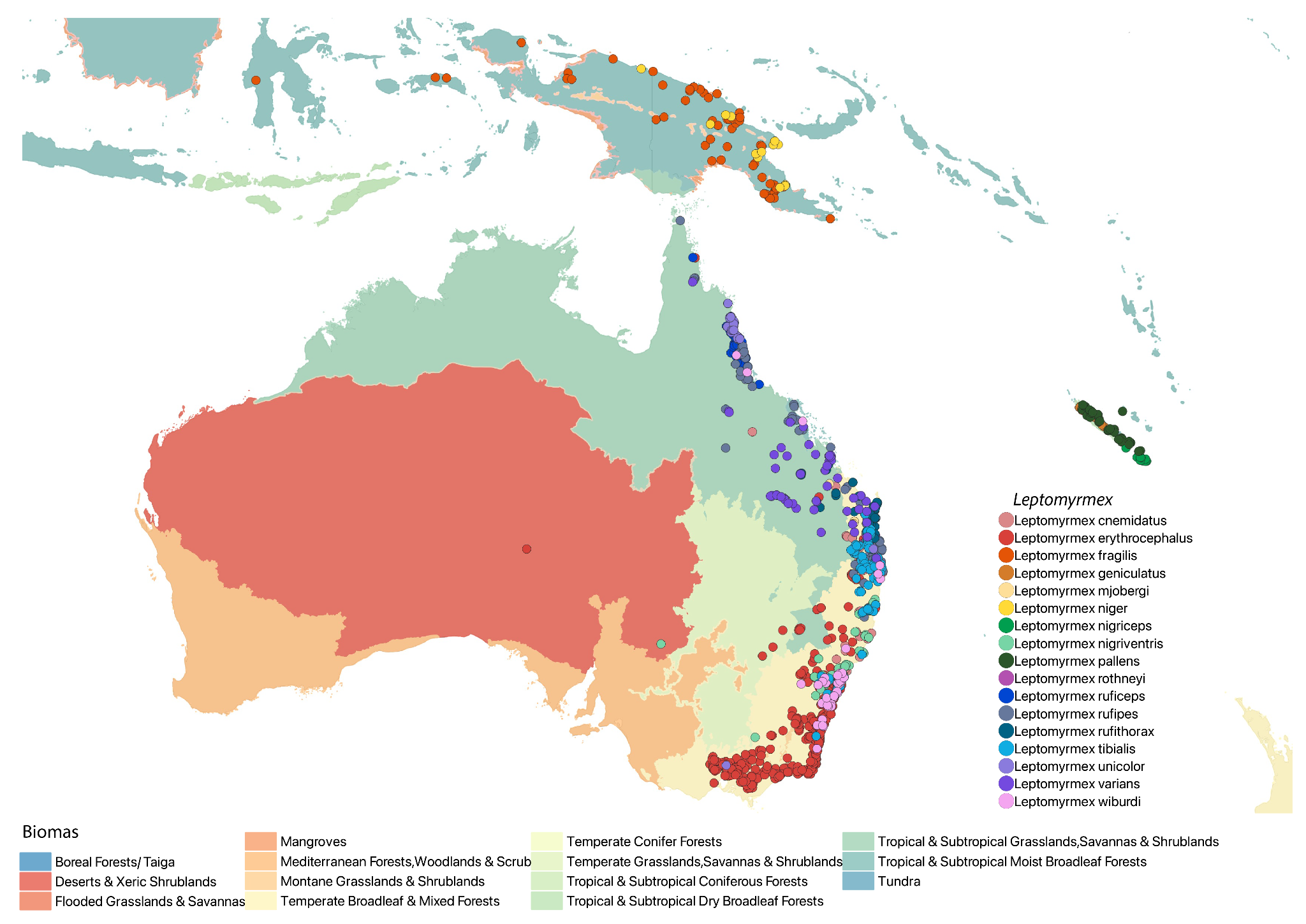

Figure S6. *Leptomyrmex* species occurrence points downloaded from GBIF.
